# Influence of elevated CO_2_ on development and food utilization of target armyworm *Mythimna separata* fed on transgenic *Bt* maize infected by nitrogen-fixing bacteria

**DOI:** 10.1101/292565

**Authors:** Zhuo Li, Yingying Song, Likun Li, Long Wang, Bin Liu, Megha N. Parajulee, Fajun Chen

## Abstract

Bt crops will face a new ecological risk of reduced effectiveness against target-insect pests owing to the general decrease in exogenous-toxin content in Bt crops grown under elevated CO_2_. How to deal with this issue may affect the sustainability of transgenic crops as an effective pest management tool especially under future CO_2_ raising. In this study, azotobacters, as being one potential biological regulator to enhance crops’ nitrogen utilization efficiency, were selected and the effects of *Bt* maize and non-*Bt* maize infected by *Azospirillum brasilense* and *Azotobacter chroococcum* on development and food utilization of target *Mythimna separate* were studied under ambient and elevated CO_2_. The results indicated that azotobacter infection significantly increased larval life-span, pupal duration, RCR and AD of *M. separata*, and significantly decreased RGR, ECD and ECI of *M. separata* fed on *Bt* maize; There were opposite trends in development and food utilization of *M. separata* fed on non-*Bt* maize infected with azotobacters compared with the buffer control regardless of CO_2_ level. Presumably, the application of azotobacter infection could make *Bt* maize facing lower field hazards from the target pest of *M. separate*, and finally improve the resistance of *Bt* maize against target lepidoptera pests especially under elevated CO_2_.

**Summary statement:** Elevated CO_2_ effect on development and food utilization of target armyworm *Mythimna separata* fed on *Bt* maize infected by azotobacter, *Azospirillum brasilense* and *Azotobacter chroococcum*

## Introduction

With increased fossil fuel combustion and drastic changes in land utilization, the concentration of atmospheric carbon dioxide (i.e., CO_2_) has increased by more than 40%, from 280 to 400 ppm, between the industrial revolution and now (Ciais et al., 2013). The recent forecast indicated that atmospheric CO_2_ concentration will increase to approximately 900 ppm by 2100 (IPCC, 2014). Increasing atmospheric CO_2_ concentration alone can be very significant in crop production because of its direct effect on plant physiology and biochemistry (Cornelissen, 2011), and indirect effect on tri-trophic interactions involving plants, herbivores, and predators or pathogens (Robinson et al., 2012; Trębicki et al., 2017). Elevated atmospheric CO_2_ also affects the crop production via direct or indirect impact on the physiology and feeding behavior of phytophagous insects (Zvereva & Kozlov, 2006; Massad & Dyer, 2010; O’Neill et al., 2010). These changes may then lead to more severe and frequent outbreaks of pest insects in agricultural ecosystems (Percy et al., 2002).

Several studies have shown that the elevated CO_2_ increased lepidopteran insect feeding and damage severity in agricultural crops (Ainsworth & Schurr, 2007; Lindroth et al., 2001), because of the increased proportion of C: N in host plant tissue and lower nutritional quality caused by elevated CO_2_ (Ainsworth & Schurr, 2007). For example, larvae of *Helicoverpa armigera* fed on wheat grown in elevated CO_2_ showed the extended larval life span and increased consumption with reduced growth rate (Chen et al., 2004). Transgenic maize that expresses insecticidal *Cry* proteins derived from the soil bacterium *Bacillus thuringiensis* Berliner (Bt) has been used to control target lepidopteran insects (Carrière et al., 2010; Huang et al., 2014; Walters et al., 2010), e.g., European corn borer *Ostrinia nubilalis* (Hübner), Asian corn borer *O. furnacalis* (Guenée) (Lepidoptera: Crambidae) and corn armyworm *Mythimna separata* (Lepidoptera: Noctuidae) (Guo et al., 2016; Zhang et al., 2010; Jia et al., 2010). Transgenic *Bt* maize has recently been adopted worldwide (Cattaneo et al., 2006; Huang et al., 2005; Hutchison et al., 2010; Lu et al., 2012). It was anticipated that the primary effect of elevated CO_2_ on *Bt* toxin production would be due to differences in N concentration in plant tissues (Coviella et al., 2002). Biologically relevant changes in plant defensive chemistry of *Bt* maize are expected to have measurable effects on the target lepidopteran pests under climate change.

Additionally, many researchers found that the nitrogen metabolism of transgenic *Bt* crops could affect the expression of *Bt* toxin protein (Stitt & Krapp, A, 1999; Pang et al., 2005; Gao et al., 2009). Nitrogen plays the most important role for plant growth, and it is an important complement of enzymes catalyzing and controlling reactions in plants for normal physiological processes (Richardson, 2009). While most of nitrogen in the environment is found in a form of nitrogen gas (N_2_) which approximately amounts to 78% in the atmosphere, plant available nitrogen found in soil is generally derived from fertilizer augmentation. As plants cannot use N_2_ directly, soil-inhabiting microbes play a significant role in nitrogen uptake by plants as they change the nitrogen gas into ammonia (Yamprai et al., 2014). *Azospirillum* sp. and *Azotobacter* sp are the two major free-living soil microbes (Biari et al., 2008; Sastro et al., 2014), that are economically important nitrogen-fixing bacteria in maize crop production system (Yamprai et al., 2014). Thus, optimization of soil-nitrogen management offers significant potential in the utilization of soil azotobacters to increase *Bt*-crop nitrogen utilization to affect the expression of *Bt* toxin under elevated CO_2_.

In this study, we examined the influence of elevated CO_2_ on larval development and food utilization of target armyworm *M. separata* fed on *Bt* maize (line IE09S034 with *Cry1Ie* gene)versus its non-*Bt* parental line (cv. Xianyu335) infected with *A. brasilense* and *A. chroococcum* under ambient and elevated CO_2_.

## Materials and Methods

### Setup of CO_2_ levels

A two-year study (2016-2017) was conducted in six open-top chambers (i.e., OTCs; Granted Patent: ZL201120042889.1; 2.5m in height × 3.2m in diameter) (Chen et al., 2011) at the Innovation Research Platforms for Climate Change, Biodiversity and Pest Management (CCBPM; http://www.ccbpm.org) field laboratory in Ningjin County, Shandong Province of China (37º38 ′ 30.7′′ N, 116º51 ′ 11.0′′ E). Two CO_2_ levels, ambient (375 μl/L, hereafter referred to as *a*CO_2_) and elevated (750 μl/L or double-ambient, hereafter referred to as *e*CO_2_) were applied continuously from 10 June to 7 October in both years. Three OTCs were used for each CO_2_ treatment, and the CO_2_ concentrations in each OTC were monitored continuously and adjusted using an infrared CO_2_ analyzer (Ventostat 8102; Telaire Company, Goleta, CA). The OTCs of elevated CO_2_ treatments were inflated with canned CO_2_ gas with 95% purity and automatically controlled by the same type of infrared CO_2_ analyzer (Chen et al., 2004). Actual mean CO_2_ concentrations (2016-*a*CO_2_: 372.6 μl/L, *e*CO_2_: 744.4 μl/L; 2017-*a*CO_2_: 374.5 μl/L, *e*CO_2_: 748.8 μl/L) and temperature (2016-26.06℃; 2017-26.11℃) throughout the entire experiment are measured in both 2016 and 2017.

### Plant materials

The *Bt* maize cultivar (Line IE09S034, hereafter referred to as Bt) and its non-*Bt* parental line (cv. Xianyu 335, referred to as Xy) were both obtained from the Institute of Crop Sciences, Chinese Academy of Agricultural Sciences. Both *Bt* and non-Bt lines used in this study possessed identical growing periods (approximately 102 d) and were well adapted to the growing conditions of northern China (Guo et al., 2016; Zhang et al., 2010; Jia et al., 2010; Ling, 2010). Both maize accessions were planted in plastic buckets (diameter × height=30 cm × 45 cm) filled with 20kg autoclaved soil and 10g compound fertilizer (N: P: K=18: 15: 12), then placed them into chambers on 10 June each year.

### Soil nitrogen-fixing bacteria and infection of maize seeds

Lyophilized *Azospirillum brasilense* (strain number ACCC 10103) and *Azotobacter chroococcum* (strain number ACCC 10006) were provided by Agriculture Culture Collection of China (ACCC) in plastic tubes (3 cm in diameter and 15 cm in height) with bacterial growth medium. Both species of azotobacters were grown in liquid medium at 28°C under continuous shaking (200rpm) until they reached an absorbance of 1.008 (*A. brasilense*) and 1.005 (*A. chroococcum*) at a wavelength of 600nm. Before inoculation, the culture was centrifuged, and the supernatant was discarded, and the pellet of cells was re-suspended in the liquid medium to a density of 10^8^ copies per milliliter. The seeds of both *Bt* and non-Bt maize were infected with *A. brasilense* and *A. chroococcum* cultures each, and the inoculation doses were all adjusted to a final volume of 10 ml. After inoculation, all the treated seeds were maintained under sterile laminar air flow for 2h at 28^˚^C (Cass°n, 2009). Bacteria inoculation treatments consisted of three types of azotobacter infection, including (1) seeds infected with *A. brasilense* (referred to as AB); (2) seeds infected with *A. chroococcum* (referred to as AC); and (3) non-infected seeds (control) treated with a final volume of buffer solution (referred to as CK). The entire experiment, thus, consisted of twelve treatments, including two CO_2_ levels (*a*CO_2_ and *e*CO_2_), two maize cultivars (Bt and Xy), and three azotobacter infections (AB, AC, and CK), replicated six times. Specifically, six buckets for each maize cultivar (Bt and Xy) and three azotobacter inoculations (6 buckets per cultivar treatment × 2 cultivars × 3 inoculation treatments=36 buckets) were placed randomly in each CO_2_ chamber (ambient and double-ambient CO_2_), and 3 maize seeds were sown in each bucket at 2 cm soil depth. No pesticides were applied during the entire experimental period and the manual weeding keep the maize buckets weed-free during the experiment. The rhizosphere soil was sampled from each bucket 1 day before planting, 14 days after planting, and at harvest and measured the relative density of AB (Sequence specific primers of *A. brasilense* NifH gene: F’-CAAGGGCACCATCCCGAC; R’-CTGCTGCTCCTCCGACT) and AC (Sequence specific primers of *A. chroococcum* nifH gene: F’- GTGACCCGAAAGCTGACTCC; R’- CCACCTTCAGCACGTCTTCC) using RT-PCR (Table 1).

**Table 1.**
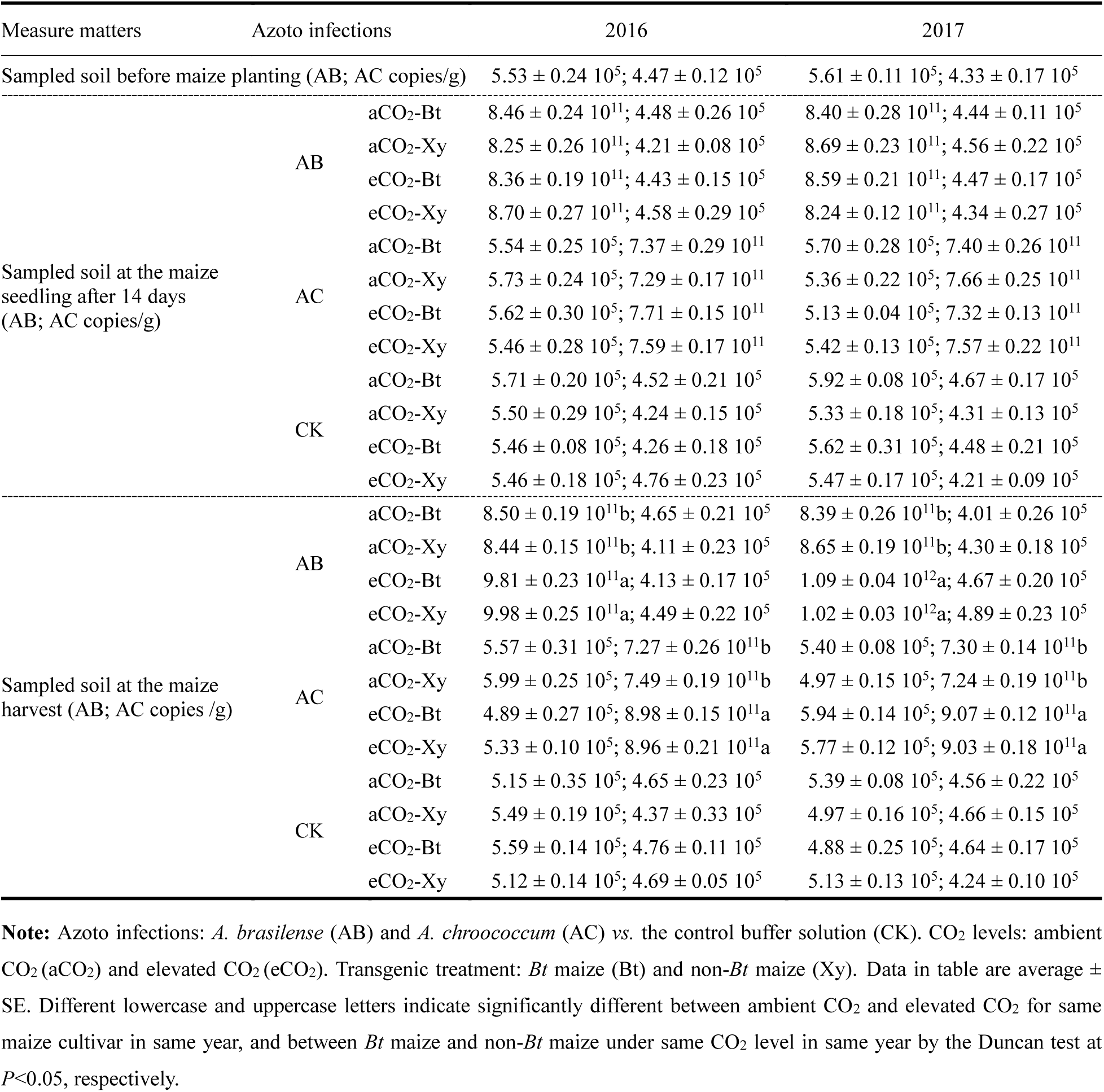
The rhizosphere soil densities of azotobacters, *Azospirillum brasilense* (i.e., AB) and *Azotobacter chroococcum*(AC) inoculated in the potted soil of transgenic *Bt* maize (i.e., Bt) and its parental line of non-*Bt* maize (Xy) grown under ambient and elevated CO_2_ in 2016 and 2017

| Measure matters | Azoto infections |  | 2016 | 2017 |
| --- | --- | --- | --- | --- |
| Sampled soil before maize planting (AB; AC copies/g) | | | $5.53 \pm 0.24 \times 10^5$; $4.47 \pm 0.12 \times 10^5$ | $5.61 \pm 0.11 \times 10^5$; $4.33 \pm 0.17 \times 10^5$ |
| Sampled soil at the maize seedling after 14 days (AB; AC copies/g) | AB | aCO<sub>2</sub>-Bt | $8.46 \pm 0.24 \times 10^{11}$; $4.48 \pm 0.26 \times 10^5$ | $8.40 \pm 0.28 \times 10^{11}$; $4.44 \pm 0.11 \times 10^5$ |
| | | aCO<sub>2</sub>-Xy | $8.25 \pm 0.26 \times 10^{11}$; $4.21 \pm 0.08 \times 10^5$ | $8.69 \pm 0.23 \times 10^{11}$; $4.56 \pm 0.22 \times 10^5$ |
| | | eCO<sub>2</sub>-Bt | $8.36 \pm 0.19 \times 10^{11}$; $4.43 \pm 0.15 \times 10^5$ | $8.59 \pm 0.21 \times 10^{11}$; $4.47 \pm 0.17 \times 10^5$ |
| | | eCO<sub>2</sub>-Xy | $8.70 \pm 0.27 \times 10^{11}$; $4.58 \pm 0.29 \times 10^5$ | $8.24 \pm 0.12 \times 10^{11}$; $4.34 \pm 0.27 \times 10^5$ |
| | AC | aCO<sub>2</sub>-Bt | $5.54 \pm 0.25 \times 10^5$; $7.37 \pm 0.29 \times 10^{11}$ | $5.70 \pm 0.28 \times 10^5$; $7.40 \pm 0.26 \times 10^{11}$ |
| | | aCO<sub>2</sub>-Xy | $5.73 \pm 0.24 \times 10^5$; $7.29 \pm 0.17 \times 10^{11}$ | $5.36 \pm 0.22 \times 10^5$; $7.66 \pm 0.25 \times 10^{11}$ |
| | | eCO<sub>2</sub>-Bt | $5.62 \pm 0.30 \times 10^5$; $7.71 \pm 0.15 \times 10^{11}$ | $5.13 \pm 0.04 \times 10^5$; $7.32 \pm 0.13 \times 10^{11}$ |
| | | eCO<sub>2</sub>-Xy | $5.46 \pm 0.28 \times 10^5$; $7.59 \pm 0.17 \times 10^{11}$ | $5.42 \pm 0.13 \times 10^5$; $7.57 \pm 0.22 \times 10^{11}$ |
| | CK | aCO<sub>2</sub>-Bt | $5.71 \pm 0.20 \times 10^5$; $4.52 \pm 0.21 \times 10^5$ | $5.92 \pm 0.08 \times 10^5$; $4.67 \pm 0.17 \times 10^5$ |
| | | aCO<sub>2</sub>-Xy | $5.50 \pm 0.29 \times 10^5$; $4.24 \pm 0.15 \times 10^5$ | $5.33 \pm 0.18 \times 10^5$; $4.31 \pm 0.13 \times 10^5$ |
| | | eCO<sub>2</sub>-Bt | $5.46 \pm 0.08 \times 10^5$; $4.26 \pm 0.18 \times 10^5$ | $5.62 \pm 0.31 \times 10^5$; $4.48 \pm 0.21 \times 10^5$ |
| | | eCO<sub>2</sub>-Xy | $5.46 \pm 0.18 \times 10^5$; $4.76 \pm 0.23 \times 10^5$ | $5.47 \pm 0.17 \times 10^5$; $4.21 \pm 0.09 \times 10^5$ |
| Sampled soil at the maize harvest (AB; AC copies /g) | AB | aCO<sub>2</sub>-Bt | $8.50 \pm 0.19 \times 10^{11}$b; $4.65 \pm 0.21 \times 10^5$ | $8.39 \pm 0.26 \times 10^{11}$b; $4.01 \pm 0.26 \times 10^5$ |
| | | aCO<sub>2</sub>-Xy | $8.44 \pm 0.15 \times 10^{11}$b; $4.11 \pm 0.23 \times 10^5$ | $8.65 \pm 0.19 \times 10^{11}$b; $4.30 \pm 0.18 \times 10^5$ |
| | | eCO<sub>2</sub>-Bt | $9.81 \pm 0.23 \times 10^{11}$a; $4.13 \pm 0.17 \times 10^5$ | $1.09 \pm 0.04 \times 10^{12}$a; $4.67 \pm 0.20 \times 10^5$ |
| | | eCO<sub>2</sub>-Xy | $9.98 \pm 0.25 \times 10^{11}$a; $4.49 \pm 0.22 \times 10^5$ | $1.02 \pm 0.03 \times 10^{12}$a; $4.89 \pm 0.23 \times 10^5$ |
| | AC | aCO<sub>2</sub>-Bt | $5.57 \pm 0.31 \times 10^5$; $7.27 \pm 0.26 \times 10^{11}$b | $5.40 \pm 0.08 \times 10^5$; $7.30 \pm 0.14 \times 10^{11}$b |
| | | aCO<sub>2</sub>-Xy | $5.99 \pm 0.25 \times 10^5$; $7.49 \pm 0.19 \times 10^{11}$b | $4.97 \pm 0.15 \times 10^5$; $7.24 \pm 0.19 \times 10^{11}$b |
| | | eCO<sub>2</sub>-Bt | $4.89 \pm 0.27 \times 10^5$; $8.98 \pm 0.15 \times 10^{11}$a | $5.94 \pm 0.14 \times 10^5$; $9.07 \pm 0.12 \times 10^{11}$a |
| | | eCO<sub>2</sub>-Xy | $5.33 \pm 0.10 \times 10^5$; $8.96 \pm 0.21 \times 10^{11}$a | $5.77 \pm 0.12 \times 10^5$; $9.03 \pm 0.18 \times 10^{11}$a |
| | CK | aCO<sub>2</sub>-Bt | $5.15 \pm 0.35 \times 10^5$; $4.65 \pm 0.23 \times 10^5$ | $5.39 \pm 0.08 \times 10^5$; $4.56 \pm 0.22 \times 10^5$ |
| | | aCO<sub>2</sub>-Xy | $5.49 \pm 0.19 \times 10^5$; $4.37 \pm 0.33 \times 10^5$ | $4.97 \pm 0.16 \times 10^5$; $4.66 \pm 0.15 \times 10^5$ |
| | | eCO<sub>2</sub>-Bt | $5.59 \pm 0.14 \times 10^5$; $4.76 \pm 0.11 \times 10^5$ | $4.88 \pm 0.25 \times 10^5$; $4.64 \pm 0.17 \times 10^5$ |
| | | eCO<sub>2</sub>-Xy | $5.12 \pm 0.14 \times 10^5$; $4.69 \pm 0.05 \times 10^5$ | $5.13 \pm 0.13 \times 10^5$; $4.24 \pm 0.10 \times 10^5$ |

**Note:** Azoto infections: *A. brasilense* (AB) and *A. chroococcum* (AC) vs. the control buffer solution (CK). CO<sub>2</sub> levels: ambient CO<sub>2</sub> (aCO<sub>2</sub>) and elevated CO<sub>2</sub> (eCO<sub>2</sub>). Transgenic treatment: *Bt* maize (Bt) and non-*Bt* maize (Xy). Data in table are average ± SE. Different lowercase and uppercase letters indicate significantly different between ambient CO<sub>2</sub> and elevated CO<sub>2</sub> for same maize cultivar in same year, and between *Bt* maize and non-*Bt* maize under same CO<sub>2</sub> level in same year by the Duncan test at *P*<0.05, respectively.

### Insect source and rearing

The colony of armyworm *M. separata* was originated from a population collected in maize fields in Kangbao County, Hebei province of China (41.87°N, 114.6°E) in the summer of 2014, and fed on artificial diet^45^ and maintained for more than 10 generations in climate-controlled growth chambers (GDN-400D-4; Ningbo Southeast Instrument CO., LTD, Ningbo, China) at 26 ± 1 °C, 65 ± 5% RH, and 14: 10h L/D photoperiod. The same rearing conditions were maintained for the following experiments. Newly-hatched larvae were randomly selected from the above colony of *M. separata* and fed on artificial diet (Bi, 1983) until the 2nd instar larvae, and then the 3rd instar larvae were individually fed on excised leaves of the experimental plants from CO_2_ chambers. Feeding trials were conducted in plastic dish (6 cm in diameter and 1.6 cm in height) and the experimental leaves were randomly selected from 6 buckets for each of the 12 experimental treatment combinations (two cultivars × two CO_2_treatments × 3 bacteria inoculations) during the heading stage until pupation. Sample size for the *M. separata* larval feeding trial consisted of 20 individuals (sample unit size) with 5 replicates for each of the 12 treatment combinations (i.e., 1200 larvae evaluated for the entire study). Because of the cannibalism among the late instar larvae of *M. separata* (Jiang et al., 2016; Ali et al., 2016; Liu et al., 2017), the sampled individuals were reared separately in the Petri dish until pupation.

### Development and reproduction of *M. separata*

Larval development was evaluated from 3rd instar to pupation by way of observing each individual petri dish every 8 h and recording the timing of larval ecdysis, pupation, and emergence of *M. separata* moths. After eclosion, the newly emerged moths were paired (female: male=1: 1) for mating in a metal frame screen cage (length × width × height=35 cm × 35 cm× 40 cm), and the paired moths were fed with a 10% honey solution provided on a large cotton wick in a single plastic cup (diameter × height=8 cm × 20 cm) covered with cotton net yarn butter paper for ovipositing. The cotton net yarn and butter paper were replaced every day. Moth survivorship and oviposition were recorded daily until both moths from each pair died.

### Food utilization of the larvae of *M. separata*

Each 3rd instar test larvae of *M. separaae* was weighed at the initiation of the feeding trial by using an electronic balance (AL104, METTLER-TOLEDO, Switzerland). Total accumulated feces from 3^rd^ instar until the larva entered pupal stage (6^th^ instar), 6th instar larval weight, and the remaining leaves were also weighed. The food utilization indices of *M. separata* included the relative growth rate (RGR), relative consumption rate (RCR), approximate digestibility (AD), efficiency of conversion of ingested food (ECI) and efficiency of conversion of digested food (ECD) (Chen et al., 2005a; Chen et al., 2005b). Formulas for calculation of the measured indices were adapted from Chen *et al.* (2005b).

### Data analysis

All data were analyzed using the statistical software SPSS 19.0 (2015, SPSS Institute, Chicago, IL). Four-way analysis of variance (ANOVA) was used to analyze the effects of CO_2_ levels (elevated *vs.* ambient), transgenic treatment (*Bt* maize *vs.* non-*Bt* maize), azoto infection (AB and AC vs. CK), sampling years (2016 *vs*. 2017), and the interactions on the measured indices of growth, development, and reproduction, including larval life-span, pupation rate, pupal weight, pupal duration, adult longevity and fecundity of *M. separata*. The measured food utilization indices were analyzed by using an analysis of covariance (ANCOVA) with initial weight of *M. separata* (i.e., 3rd instar larva) as a covariate for RCR and RGR, while food consumption was a covariate for ECI and AD to correct the effect of variation in the growth and food assimilation of *M. separata* (Raubenheimer & Simpson, 1992); food assimilated was also used as a covariate to analyze the ECD parameter (Hägele, 1999). The assumption of a parallel slope between covariate and dependent variable was satisfied for each analysis. Treatment means were separated by using the Duncan-test to examine significant difference at *P*<0.05.

## Results

### Effects of CO_2_ level, transgenic treatment and azoto infection on the rhizosphere soil densities of *A. brasilense* and *A. chroococcum* in different sampling period

Significant effects of azoto infection (*P*<0.001) were observed on the measured rhizosphere soil densities of *A. brasilense* (AB) and *A. chroococcum* (AC) 14 days after maize planting. Compared with ambient CO_2_, elevated CO_2_ significantly increased the rhizosphere soil densities of both *A. brasilense* and *A. chroococcum*; compared with the control buffer solution (CK), azoto infection significantly increased the rhizosphere soil densities of *A. brasilense* and *A. chroococcum* (*P*<0.001; Table 1). CO_2_ level and azoto infection (*P*<0.001) both significantly affected the measured rhizosphere soil densities of *A. brasilense* and *A. chroococcum* at maize harvest (Table 2).

**Table 2.** Four-way ANOVA on the rhizosphere soil densities of *Azospirillum brasilense* (AB) and *Azotobacter chroococcum* (AC) inoculated in the potted soil of *Bt* maize and its parental line of non-*Bt* maize treatments grown under ambient and elevated CO_2_ in 2016 and 2017 (F/P values)

| Impact factors | Sampled soil at the maize seedling after 14 days |  | Sampled soil at the maize harvest |  |
| --- | --- | --- | --- | --- |
|  | AB | AC | AB | AC |
| Y<sup>a</sup> | 0.00/0.99 | 0.002/0.96 | 0.97/0.33 | 1.13/0.29 |
| Cv.<sup>b</sup> | 0.32/0.574 | 0.36/0.55 | 0.070/0.79 | 0.080/0.78 |
| CO<sub>2</sub><sup>c</sup> | 0.19/0.89 | 0.80/0.38 | 26.01/<0.001*** | 331.16/<0.001*** |
| Azoto<sup>d</sup> | 2555.00/<0.001*** | 1380.37/<0.001*** | 1311.83/<0.001*** | 2080.71/<0.001*** |
| Y × Cv. | 0.62/0.44 | 1.96/0.17 | 0.18/0.673 | 0.53/0.47 |
| Y × CO<sub>2</sub> | 1.21/0.28 | 2.50/0.12 | 0.74/0.40 | 0.032/0.86 |
| Y × Azoto | 0.00/1.00 | 0.02/0.99 | 0.97/0.39 | 1.13/0.33 |
| Cv. × CO<sub>2</sub> | 0.35/0.55 | 0.010/0.92 | 0.32/0.57 | 0.36/0.55 |
| Cv. × Azoto | 0.32/0.73 | 0.36/0.70 | 0.070/0.93 | 0.80/0.93 |
| CO<sub>2</sub> × Azoto | 0.019/0.98 | 0.80/0.45 | 26.01/<0.001*** | 331.16/<0.001*** |
| Y × Cv. × CO<sub>2</sub> | 5.99/0.018* | 0.06/0.94 | 0.82/0.37 | 0.63/0.43 |
| Y × Cv. × Azoto | 0.62/0.54 | 1.96/0.15 | 0.18/0.84 | 0.53/0.59 |
| Y × CO<sub>2</sub> × Azoto | 1.21/0.31 | 2.50/0.093 | 0.74/0.49 | 0.032/0.97 |
| Cv. × CO<sub>2</sub> × Azoto | 0.35/0.70 | 0.10/0.99 | 0.32/0.73 | 0.36/0.70 |
| Y × Cv. × CO<sub>2</sub> × Azoto | 5.99/0.05 | 0.006/0.99 | 0.82/0.45 | 0.63/0.54 |

**Note:** \* $P < 0.05$ , \*\* $P < 0.01$ , \*\*\* $P < 0.001$ ; <sup>a</sup>: Year (2016 vs. 2017); <sup>b</sup>: Transgenic treatment (*Bt* maize vs. non-*Bt* maize); <sup>c</sup>: CO<sub>2</sub> levels (elevated CO<sub>2</sub> vs. ambient CO<sub>2</sub>); <sup>d</sup>: Azoto infection (*A. brasilense* and *A. chroococcum* vs. the buffer control).

### Effects of CO_2_ level, transgenic treatment and azoto infection on the development and reproduction of *M. separata*

CO_2_ level and transgenic treatment both significantly affected the larval life-span, pupation rate, pupal weight and duration, adult longevity, and fecundity in *M. separata* fed on both *Bt* and non-*Bt* maize infected with *A. brasilense* and *A. chroococcum* (*P*<0.001). However, the azoto infection significantly affected the larval life-span, pupal duration (*P*<0.05) and fecundity (*P*<0.001) of *M. separata* fed on both cultivars and at both CO_2_ levels (Table 3).

**Table 3.** Four-way ANCOVA on the growth, development, reproduction and food utilization indices of armyworm, *Mythimna separata* fed on *Bt* maize (i.e., Bt) and its parental line of non-*Bt* maize (Xy) during the heading stage, infected with azotobacters (Azoto), *A. brasilense* (i.e., AB) and *A. chroococcum* (AC) as well buffer solution as the control (CK) under ambient and elevated CO_2_ in 2016 and 2017 (*F/P* values)

| Impact factors | Larval life-span<br>(day (n=828)) | Pupation rate<br>(% (n=828)) | Pupal weight<br>(g (n=663)) | Pupal duration<br>(day (n=663)) | Adult longevity<br>(day (n=576)) | Fecundity<br>(eggs per female<br>(n=198)) | The 3<sup>rd</sup>- 6<sup>th</sup> instar larvae (n=828) |  |  |  |  |
| --- | --- | --- | --- | --- | --- | --- | --- | --- | --- | --- | --- |
|  |  |  |  |  |  |  | RGR<br>(mg g<sup>-1</sup>day<sup>-1</sup>) | RCR<br>(mg g<sup>-1</sup>day<sup>-1</sup>) | AD<br>(%) | ECD<br>(%) | ECI<br>(%) |
| Covariate<sup>e</sup> |  |  |  |  |  |  | 1.81/0.11 | 3.83/0.067 | 0.781/0.23 | 0.580/0.41 | 0.87/0.32 |
| Y<sup>a</sup> | 1.62/0.32 | 0.94/0.71 | 1.21/0.62 | 0.11/0.90 | 1.65/0.33 | 2.99/0.10 | 1.19/0.31 | 3.96/0.12 | 4.56/0.061 | 4.82/0.059 | 5.80/0.053 |
| Cv.<sup>b</sup> | 1662.03/<0.001*** | 329.16/<0.001*** | 275.04/<0.001*** | 229.78/<0.001*** | 450.27/<0.001*** | 2032.99/<0.001*** | 1545.53/<0.001*** | 302.67/0.<0.001*** | 185.62/0.<0.001*** | 716.17/0.<0.001*** | 1038.95/0.<0.001*** |
| CO<sub>2</sub><sup>c</sup> | 62.13/<0.001*** | 55.53/<0.001*** | 12.44/0.001** | 19.02/<0.001*** | 41.62/<0.001*** | 278.70/<0.001*** | 67.09/<0.001*** | 27.98/<0.001*** | 9.69/0.003** | 35.90/<0.001*** | 57.98/<0.001*** |
| Azoto<sup>d</sup> | 3.64/0.034* | 2.53/0.077 | 1.20/0.63 | 7.04/0.011* | 0.35/0.70 | 27.54/<0.001*** | 12.26/<0.001*** | 26.84/<0.001*** | 35.28/<0.001*** | 13.64/<0.001*** | 7.22/0.002** |
| Y × Cv. | 9.22/0.004** | 2.22/0.14 | 2.22/0.14 | 1.98/0.32 | 3.10/0.054 | 0.102/0.75 | 4.27/0.049* | 5.69/0.021* | 1.86/0.18 | 19.21/<0.001*** | 15.64/<0.001*** |
| Y × CO<sub>2</sub> | 5.33/0.025* | 2.16/0.20 | 0.067/0.80 | 0.17/0.85 | 13.53/<0.001*** | 2.74/0.10 | 6.90/0.012* | 4.82/0.033* | 6.73/0.013* | 6.22/0.016* | 3.41/0.071 |
| Y × Azoto | 3.63/0.034* | 1.69/0.058 | 0.63/0.57 | 0.48/0.71 | 0.59/0.64 | 1.87/0.053 | 5.04/0.010* | 0.17/0.84 | 0.27/0.77 | 0.436/0.65 | 0.25/0.78 |
| Cv. × CO<sub>2</sub> | 19.11/<0.001*** | 6.62/<0.001*** | 2.68/0.008** | 14.11/<0.001*** | 8.26/0.006** | 5.86/0.049* | 30.44/<0.001*** | 7.39/0.009** | 1.40/0.043* | 1.80/0.017* | 1.61/0.011* |
| Cv. × Azoto | 224.53/<0.001*** | 73.67/<0.001*** | 30.41/<0.001*** | 42.06/<0.001** | 26.38/<0.001*** | 195.08/<0.001*** | 213.46/<0.001*** | 48.31/<0.001*** | 73.42/<0.001*** | 132.82/<0.001*** | 144.91/<0.001*** |
| CO<sub>2</sub> × Azoto | 12.81/<0.001*** | 12.22/0.004** | 11.63/0.005** | 14.98/<0.001*** | 4.24/0.02* | 22.02/<0.001*** | 13.25/<0.001*** | 6.70/0.003** | 9.78/<0.001*** | 13.01/<0.001*** | 12.63/<0.001*** |
| Y × Cv. × CO<sub>2</sub> | 0.01/0.92 | 1.07/0.55 | 0.10/0.75 | 1.16/0.52 | 0.006/0.94 | 4.25/0.085 | 0.220/0.64 | 1.83/0.18 | 4.16/0.047* | 0.743/0.39 | 0.56/0.46 |
| Y × Cv. × Azoto | 1.55/0.33 | 1.53/0.17 | 0.064/0.94 | 2.02/0.23 | 0.46/0.68 | 8.87/0.071 | 3.27/0.047* | 0.55/0.58 | 2.23/0.12 | 4.01/0.025* | 1.41/0.25 |
| Y × CO<sub>2</sub> × Azoto | 0.063/0.69 | 0.51/0.76 | 0.48/0.62 | 1.72/0.45 | 1.78/0.18 | 14.61/0.045* | 0.72/0.49 | 1.88/0.16 | 1.33/0.28 | 2.66/0.080 | 2.47/0.095 |
| Cv. × CO<sub>2</sub> × Azoto | 8.88/0.006** | 12.61/0.003** | 7.41/0.011* | 5.66/0.005** | 13.55/0.002** | 24.04/0.000*** | 0.62/0.043* | 13.24/<0.001*** | 9.84/<0.001*** | 9.59/<0.001*** | 9.92/<0.001*** |
| Y × Cv. × CO<sub>2</sub> × Azoto | 0.19/0.83 | 0.33/0.71 | 0.14/0.87 | 0.32/0.72 | 1.73/0.30 | 0.36/0.70 | 0.48/0.62 | 0.087/0.92 | 0.98/0.38 | 0.59/0.56 | 0.06/0.94 |

**Note:** <sup>e</sup> Initial weight as a covariate for RGR and RCR, and food consumption as a covariate for AD and ECI, and food assimilated as a covariate for ECD.

Compared with ambient CO_2_, elevated CO_2_ significantly prolonged the larval life-span (+6.21%), pupal duration (+5.56%), and significantly decreased the pupation rate (-18.08%), pupal weight (-8.12%), adult longevity (-6.06%) and fecundity (-22.58%) of *M. separata* (*P*<0.05; Table 4). Also, compared with the buffer control, azoto infection with *A. brasilense* and *A. chroococcum* both significantly shortened the larval life-span (-5.20% and -5.70%), pupal duration (-3.68% and -3.81%) and fecundity (-10.20% and -9.53%) of *M. separata* (*P*<0.05; Table 4). Moreover, *Bt* maize significantly prolonged the larval life-span (+13.67%) and pupal duration (+7.54%), shortened the adult longevity (-10.41%), and decreased the pupation rate (-75.55%), pupal weight (-13.54%) and fecundity (-75.46%) of *M. separata* compared to that for non-Bt maize (*P*<0.05; Table 4).

**Table 4.** The growth, development, reproduction and food utilization of armyworm, *M. separata* fed on the detached leaves of *Bt* maize (i.e., Bt) and its parental line of non-*Bt* maize (Xy) during the heading stage, infected with azotobacters (Azoto), *A. brasilense* (i.e., AB) and *A. chroococcum* (AC) as well buffer solution as the control (CK) under ambient and elevated CO_2_ in 2016 and 2017

| Impact factors | Factor levels | Larval life-span<br>(day) | Pupation rate<br>(%) | Pupal weight<br>(g) | Pupal duration<br>(day) | Adult longevity<br>(day) | Fecundity<br>(eggs per female) | The 3<sup>rd</sup>- 6<sup>th</sup> instar larvae |  |  |  |  |
| --- | --- | --- | --- | --- | --- | --- | --- | --- | --- | --- | --- | --- |
|  |  |  |  |  |  |  |  | RGR<br>(mg g<sup>-1</sup>day<sup>-1</sup>) | RCR<br>(mg g<sup>-1</sup>day<sup>-1</sup>) | AD (%) | ECD (%) | ECI (%) |
| Cv. | Bt | 24.19±0.16a | 40.98±2.45b | 0.192±0.007b | 10.98±0.18a | 6.63±0.14b | 171.50±13.27b | 82.79±6.43b | 1636.31±13.23a | 56.13±0.72a | 9.26±0.98b | 5.13±0.57b |
|  | Xy | 21.28±0.17b | 71.94±2.38a | 0.218±0.006a | 10.21±0.19b | 7.32±0.09a | 300.92±28.64a | 94.26±7.16a | 1403.36±14.80b | 52.03±0.69b | 13.08±1.23a | 6.77±0.66a |
| CO<sub>2</sub> | Elevated | 23.42±0.20a | 51.78±2.69b | 0.197±0.007b | 10.78±0.18a | 6.77±0.11b | 212.25±19.13b | 84.33±8.67b | 1595.24±16.14a | 55.55±0.87a | 10.34±1.52b | 5.46±0.76b |
|  | Ambient | 22.05±0.17b | 61.14±2.68a | 0.213±0.005a | 10.41±0.19b | 7.18±0.13a | 260.17±21.87a | 92.72±6.59a | 1444.43±15.01b | 52.61±0.81b | 12.00±1.21a | 6.44±0.63a |
| Azoto infection | AB | 22.53±0.16b | 56.06±2.24 | 0.205±0.004 | 10.54±0.09b | 6.97±0.24 | 228.00±15.27b | 89.64±5.52a | 1549.07±9.32a | 55.06±0.85a | 10.78±1.35b | 5.75±0.67b |
|  | AC | 22.42±0.15b | 55.53±2.01 | 0.204±0.004 | 10.53±0.08b | 6.96±0.24 | 229.38±17.46b | 90.34±5.52a | 1559.90±9.32a | 54.88±0.85a | 10.96±1.35b | 5.82±0.67b |
|  | CK | 23.25±0.16a | 57.79±2.58 | 0.206±0.005 | 10.72±0.13a | 6.99±0.18 | 251.25±23.21a | 85.58±6.14b | 1450.73±9.54b | 52.30±0.62b | 11.78±1.13a | 6.28±0.66a |

**Note:** Data in table are average ± SE. Different lowercase letters indicate significant difference between treatments by the Duncan's test at $P<0.05$ .

### Impacts of CO_2_ level, transgenic treatment and azoto infection on the food utilization of *M. separata*

There were significant effects of CO_2_ level, transgenic treatment, and azoto infection (*P*<0.01 or *P*<0.001) on food utilization of *M. separata* fed on both *Bt* and non-*Bt* maize infected with *A. brasilense* and *A. chroococcum* at both CO_2_ levels in both years of the study (Table 3).

Compared with ambient CO_2_, elevated CO_2_ significantly reduced the RGR (-9.95%), ECD (-16.05%) and ECI (-17.95%), and significantly enhanced the RCR (+10.44%) and AD (+5.59%) of *M. separata* (*P*<0.05; Table 4). Compared with the buffer control, azoto infection with *A. brasilense* and *A. chroococcum* both significantly decreased the ECD (-9.28% and -7.48%) and ECI (-9.22% and -7.91%), and significantly increased the RGR (+4.75% and +5.56%), RCR (+6.78% and +7.53%) and AD (+5.28% and +4.93%) in *M. separata* (*P*<0.01; Table 4). Moreover, significant decreases in RGR (-13.85%), ECD (-41.25%) and ECI (-31.97%), and significant increases in RCR (+16.60%) and AD (+7.88%) were found when *M. separata* fed on *Bt* maize compared to that on non-*Bt* maize (*P*<0.05; Table 4).

### Interactive influence of CO_2_ level, transgenic treatment, and azoto infection on growth, development and reproduction of *M. separata*

In addition to the significant main effects of CO_2_ level, transgenic treatment, and azoto infection, there were significant two-way and three-way interaction of these three main effects on larval life-span, pupation rate, pupal weight and duration, adult longevity, and fecundity of *M. separata* fed on *Bt* and non-*Bt* maize infected with *A. brasilense* and *A. chroococcum* under both CO_2_ levels in both years of the study (*P*<0.05, *P*<0.01 or *P*<0.001; Table 3).

### Cultivar × CO_2_

Similar trends were found in the measured growth, development and reproduction indexes of *M. separata* fed on both *Bt* and non-*Bt* maize cultivars grown under elevated CO_2_ in contrast to ambient CO_2_, infected with *A. brasilense* (AB) and *A. chroococcum* (AC) as well as the control buffer (CK) in 2016 and 2017 (Fig. 1 A-F). Compared with ambient CO_2_, elevated CO_2_ significantly prolonged the larval life-span (*Bt* mazie: +6.44%; non-*Bt* maize: +8.39%) and pupal duration (non-*Bt* maize: +7.27%) and shortened the adult longevity (non-*Bt* maize: -6.19%), and significantly decreased the pupation rate (non-*Bt* maize:-20.81%), pupal weight (*Bt* mazie: -7.03%; non-*Bt* maize: -13.73%) and fecundity (*Bt* maize: -29.43%; non-*Bt* maize: -18.85%) when *M. separata* fed on *Bt* maize and non-*Bt* maize (*P*<0.05; Fig. 1A-F).

**Fig 1.**
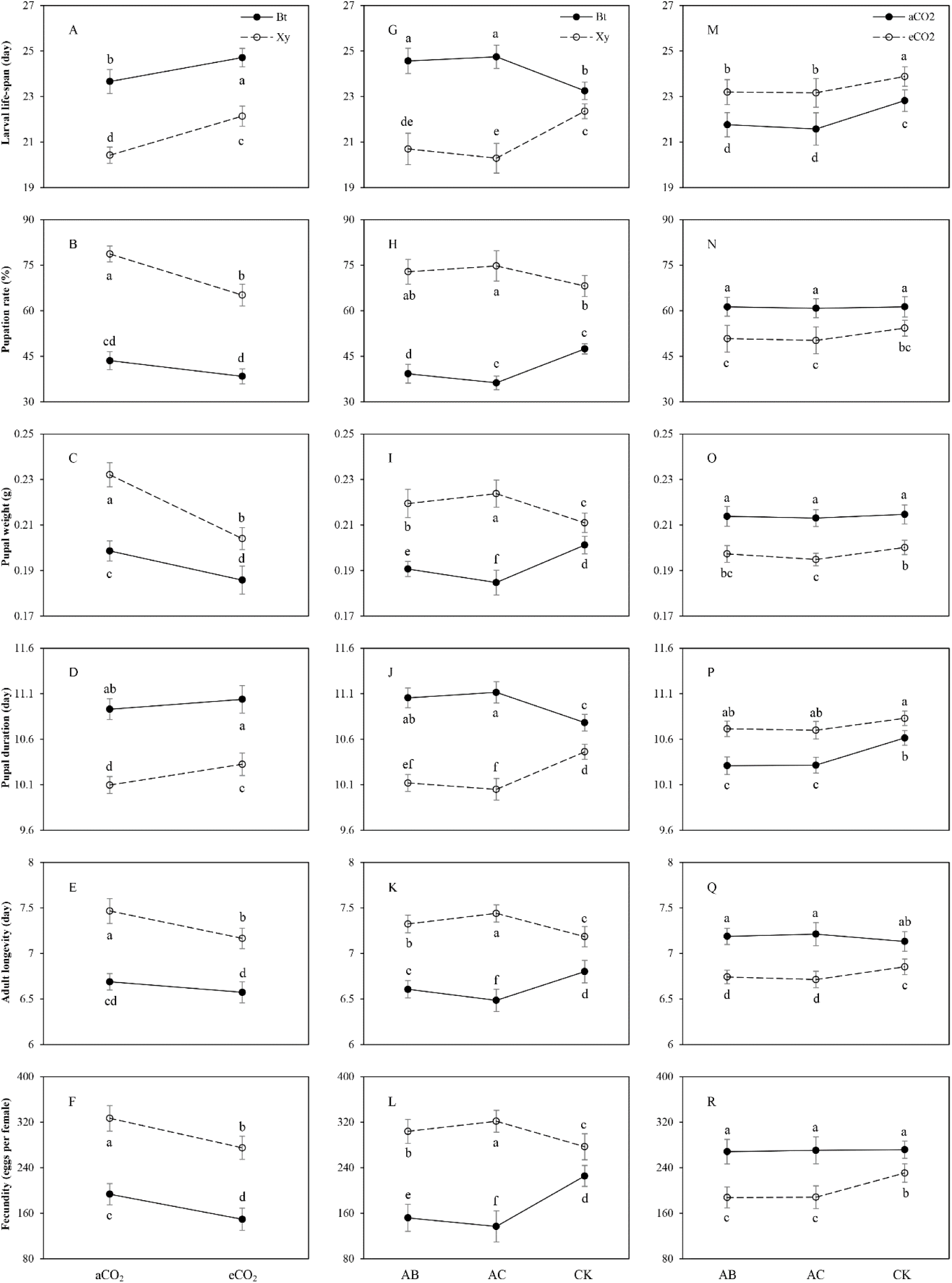
Effects of the bi-interactions between transgenic treatment and CO_2_ level (A–F), between transgenic treatment and azoto infection (G–L) and between CO_2_ level and azoto infection (M–R) on the growth, development and reproduction of armyworm, *Mythimna separata* fed on the detached leaves of *Bt* maize (i.e., Bt) and its parental line of non-*Bt* maize (Xy) during the heading stage, infected with *Azospirillum brasilense* (i.e., AB) and *Azotobacter chroococcum* (AC) as well buffer solution as the control (CK) under ambient and elevated CO_2_ in 2016 and 2017. Larval life-span–A, G, M; Pupation rate–B, H, N; Pupal weight–C, I, O; Pupal duration–D, J, P; Adult longevity–E, K, Q; Fecundity–F, L, R; Each value represents the average (±*SE*). Different lowercase letters indicate significant differences treatments by the Duncan test at *P*<0.05.

### Cultivar × Azoto

Inverse trend was found in the measured growth, development and reproduction indexes of *M. separata* fed on *Bt* maize and non-*Bt* maize, which were infected with *A. brasilense* (AB) and *A. chroococcum* (AC) under ambient and elevated CO_2_ in 2016 and 2017 (Fig. 1 G-L). Compared with the buffer control (CK), azoto infection significantly prolonged the larval life-span (AB: +7.63%; AC: +8.45%), pupal duration (AB: +4.53%; AC: +5.08%) and shortened the adult longevity (AB: -4.88%; AC: -6.94%), and decreased pupation rate (AB: -20.83%; AC: -30.81%), pupal weight (AB: -7.24%; AC: -10.65%) and fecundity (AB: -48.36%; AC: -64.60%) when *M. separata* larvae fed on *Bt* maize (*P*<0.01; Fig.1 G-L); and azoto infection significantly shortened the larval life-span (AB: -7.97%; AC: -10.15%), pupal duration (AB: -5.36%; AC: -6.08%) and prolonged the adult longevity (AB: +3.95%; AC: +5.62%), and increased pupation rate (AC: +9.73%), pupal weight (AB: +6.27%; AC: +8.16%) and fecundity (AB: +9.75%; AC: +16.16%) when *M. separata* larvae fed on non-*Bt* maize (*P*<0.01; Fig.1 G-L).

### CO_2_ × Azoto

Similar trends were found in the larval life-span, pupation rate, pupal weight and pupal duration, while inverse trends were observed in adult longevity and fecundity of *M. separata* under ambient and elevated CO_2_, which fed on *Bt* maize versus non-*Bt* maize infected with *A. brasilense* (AB) and *A. chroococcum* (AC) as well as the control buffer (CK) in 2016 and 2017 (Fig. 1 M-R). Compared with the buffer control (CK), azoto infection significantly shortened the larval life-span (AB: -6.92%; AC: -7.84%) and pupal duration (AB: -5.01%; AC: -5.01%) of *M. separata* under ambient CO_2_, and significantly shortened the larval life-span (AB: -4.98%; AC: -5.11%) and decreased the pupal weight (AC: -4.77%) under elevated CO_2_ (*P*<0.05; Fig. 1 M-P); and azoto infection significantly decreased the adult longevity (AB: -4.63%; AC: -5.09%) and fecundity (AB: -22.90%; AC: -22.58%) of *M. separata* under elevated CO_2_ (*P*<0.05; Fig. 1 Q and R).

### Cultivar × CO_2_ × Azoto

There were opposite trends in the measured growth, development and reproduction indexes of *M. separata* fed on *Bt* maize and non-*Bt* maize infected with *A. brasilense* (AB) and *A. chroococcum* (AC) compared with the buffer control (CK) in 2016 and 2017 regardless of CO_2_ level (Fig.2). In comparison with the buffer control, azoto infection with *A. brasilense* and *A. chroococcum* both significantly prolonged the larval life-span and pupal duration of *M. separata* fed on *Bt* maize, and significantly shortened the larval life-span and pupal duration of *M. separata* fed on non-*Bt* maize under the same CO_2_ level; and azoto infection with *A. brasilense* and *A. chroococcum* both significantly reduced the pupation rate, pupal weight, adult longevity and fecundity of *M. separata* fed on *Bt* maize, and significantly enhanced the pupation rate, pupal weight, adult longevity and fecundity of *M. separata* fed on non-*Bt* maize under the same CO_2_ level. Moreover, compared with ambient CO_2_, there were opposite trends in the larval life-span, pupal weight, pupal duration, adult longevity and fecundity of *M. separata* fed on *Bt* maize infected with *A. brasilense* and *A. chroococcum* compared with the buffer control under elevated CO_2_ in both years; compared with ambient CO_2_, elevated CO_2_ significantly decreased the pupation rate of *M. separata* fed on *Bt* maize, and decreased the pupation rate, pupal weight, adult longevity and fecundity of *M. separata* fed on non-*Bt* maize, and prolonged the larval life-span and pupal duration of *M. separata* fed on non-*Bt* maize infected with *A. brasilense* and *A. chroococcum* compared with the buffer control in both years.

**Fig 2.**
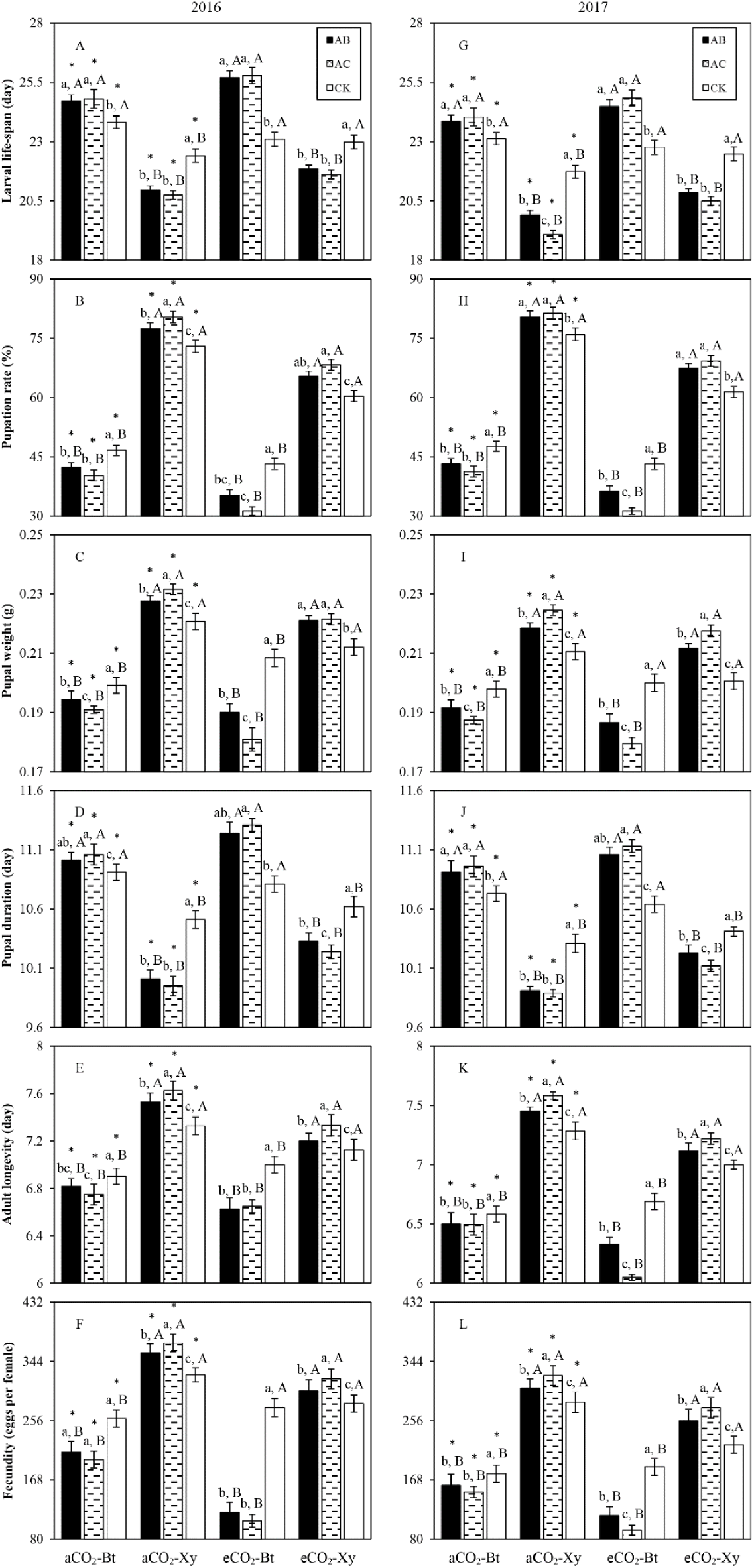
Impacts of the tri-interactions among CO_2_ level, transgenic treatment and azoto infection on the growth, development and reproduction of *M. separata* fed on the detached leaves of *Bt* maize (i.e., Bt) and non-*Bt* maize (Xy) during the heading stage, infected with *A. brasilense* (i.e., AB) and *A. chroococcum* (AC) as well buffer solution as the control (CK) under ambient and elevated CO_2_ in 2016 (A–F) and 2017 (G–L). Each value represents the average (+*SE*). Different lowercase and uppercase letters, and * indicated significant difference among three types of azoto infection for same type of maize under same CO_2_ level, between *Bt* maize and non-Bt maize for same type of azoto infection under same CO_2_ level, and between ambient and elevated CO_2_ for same type of maize and azoto infection by the Duncan test at *P*<0.05 respectively.

### Interactive effects of CO_2_ level, transgenic cultivar treatment, and azotobacter infection on food utilization of *M. separata*

In addition to significant main effects of CO_2_ level, transgenic cultivar, and azoto infection, two- and three-way interactions of these factors influenced the RGR, RCR, AD, ECD and ECI of *M. separata* larvae fed on *Bt* maize and non-*Bt* maize infected with *A. brasilense* and *A. chroococcum* under ambient and elevated CO_2_ in both years (*P*<0.05, *P*<0.01 or *P*<0.001; Table 3).

### Cultivar × CO_2_

Similar trends were found in the measured food utilization indexes of *M. separata* fed on *Bt* maize (Bt) and non-*Bt* maize (Xy) grown under elevated CO_2_ in contrast to ambient CO_2_, infected with *A. brasilense* (AB) and *A. chroococcum* (AC) as well as the control buffer (CK) in 2016 and 2017 (Fig. 3 A-E). Compared with ambient CO_2_, elevated CO_2_ significantly decreased the RGR (non-*Bt* maize: -7.34%), ECD (*Bt* mazie: -9.67%; non-*Bt* maize: -10.25%) and ECI (*Bt* maize: -8.53%; non-*Bt* maize: -8.89%), and significantly increased the RCR (*Bt* mazie: +9.69%; non-*Bt* maize: +6.37%) when *M. separata* larvae fed on *Bt* maize and non-*Bt* maize (*P*<0.05; Fig. 3 A-E).

**Fig 3.**
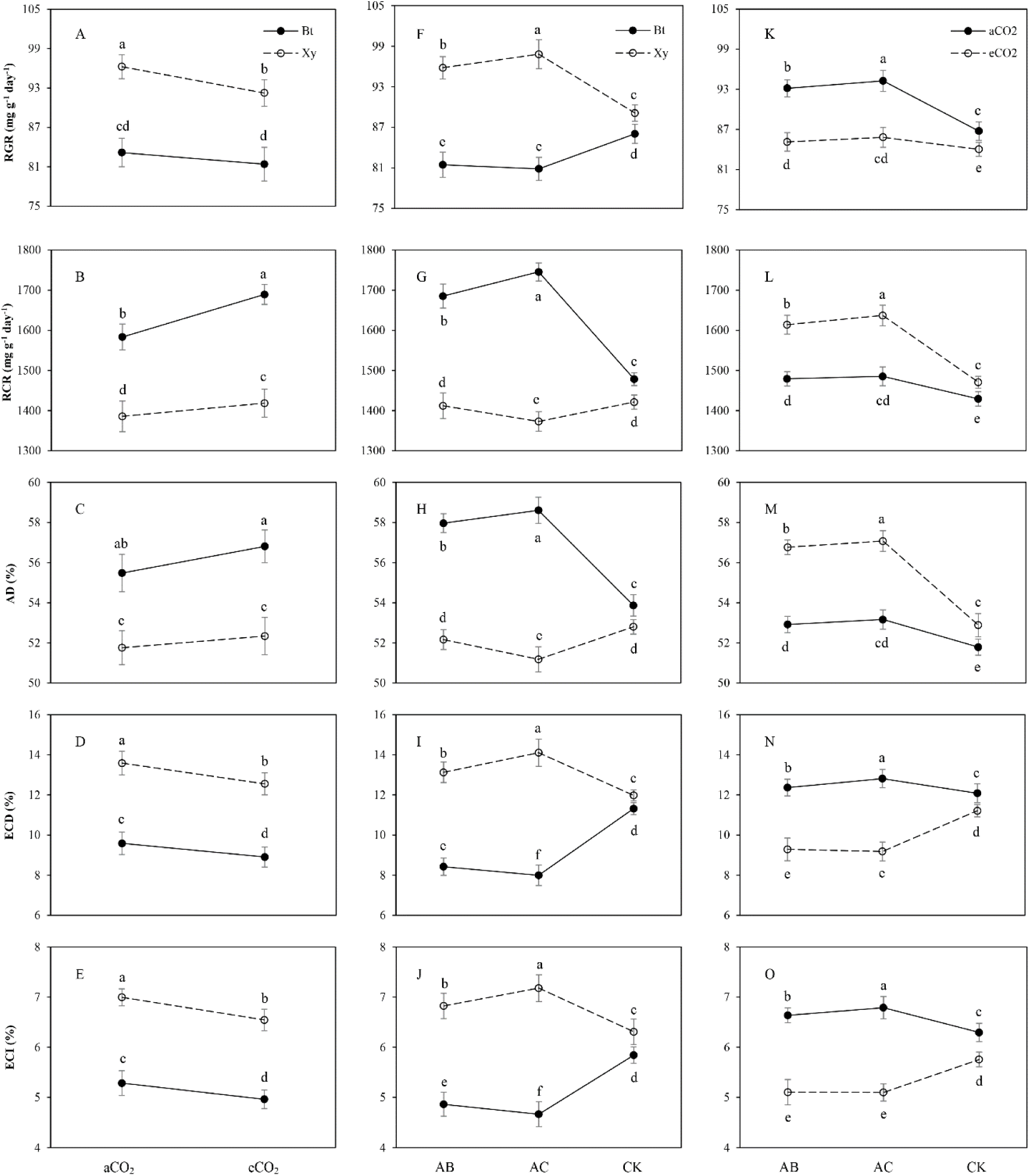
Effects of the bi-interactions between transgenic treatment and CO_2_ level (A–E), between transgenic treatment and azoto infection (F–J) and between CO_2_ level and azoto infection (K–O) on the food utilization of armyworm, *Mythimna separata* from the 3rd to the 6th instar larvae fed on the detached leaves of *Bt* maize (i.e., Bt) and its parental line of non-*Bt* maize (Xy) during the heading stage, infected with *Azospirillum brasilense* (i.e., AB) and *Azotobacter chroococcum* (AC) as well buffer solution as the control (CK) under ambient and elevated CO_2_ in 2016 and 2017. RGR–A, F, K; RCR–B, G, L; AD–C, H, M; ECD–D, I, N; ECI–E, J, O; Each value represents the average (±*SE*). Different lowercase letters indicate significant differences treatments by the Duncan test at *P*<0.05.

### Cultivar × Azoto

Inverse trend was found in the measured food utilization indexes of *M. separata* fed on *Bt* maize and non-*Bt* maize, which were infected with *A. brasilense* (AB) and *A. chroococcum* (AC) under ambient and elevated CO_2_ in 2016 and 2017 (Fig. 3 F-J). Compared with the buffer control (CK), azoto infection significantly enhanced the RGR (AB: +9.53%; AC: +11.78%), ECD (AB: +11.61%; AC: +19.79%) and ECI (AB: +10.08%; AC: +15.79%), and significantly decreased the RCR (AC: -6.52%) and AD (AC: -6.19%) when *M. separata* larvae fed on non-*Bt* maize (*P*<0.001; Fig. 3 F-J); and azoto infection significantly decreased the RGR (AB: -9.62%; AC: -10.41%), ECD (AB: -34.32%; AC: -41.55%) and ECI (AB: -20.16%; AC: -25.28%), and significantly increased the RCR (AB: +14.99%; AC: +19.06%) and AD (AB: +9.60%; AC: +10.79%) when *M. separata* larvae fed on *Bt* maize (*P*<0.001; Fig.3 F-J).

### CO_2_ × Azoto

Similar trends were observed in RGR, RCR and AD, while inverse trends were shown in ECD and ECI of *M. separata* under ambient and elevated CO_2_, which fed on *Bt* maize and non-*Bt* maize infected with *A.brasilense* (AB) and *A. chroococcum* (AC) versus control buffer (CK) (Fig. 3 K-O). Compared with the buffer control (CK), azoto infection significantly decreased ECD (AB: -20.71%; AC: -22.07%) and ECI (AB: -12.77%; AC: -12.89%) of *M. separata* larvae under elevated CO_2_, and significantly increased ECD (AB: +5.35%; AC: +8.04%) and ECI (AB: +7.43%; AC: +9.85%) of *M. separata* larvae under ambient CO_2_ (*P*<0.05; Fig.3); and azoto infection significantly enhanced RGR (AB: +3.32% and +7.40%; AC: +5.14% and +8.67%), RCR (AB: +9.78% and +5.29%,; AC: +11.32% and +5.93%) and AD (AB: +7.34% and +4.18%; AC: +7.92% and +4.66%) under elevated and ambient CO_2_, respectively (*P*<0.01; Fig. 3 K-O).

### Cultivar × CO_2_ × Azoto

There were opposite trends in the measured food utilization indexes of *M. separata* larvae fed on *Bt* maize (Bt) and non-*Bt* maize infected with *A. brasilense* (AB) and *A. chroococcum* (AC) compared with the buffer control (CK) in both years regardless of CO_2_ level (Fig.4). In comparison with the buffer control, azoto infection with *A. brasilense* and *A. chroococcum* both significantly decreased RGR, ECD and ECI of *M. separata* fed on *Bt* maize, and significantly increased RGR, ECD and ECI of *M. separata* fed on non-*Bt* maize under the same CO_2_ level; and azoto infection with *A. brasilense* and *A. chroococcum* both significantly enhanced RCR and AD of *M. separata* fed on *Bt* maize, and significantly reduced RCR and AD of *M. separata* larvae fed on non-*Bt* maize under the same CO_2_ level. Moreover, compared with ambient CO_2_, elevated CO_2_ significantly increased RCR and AD, and significantly decreased RGR, ECD, and ECI of *M. separata* larvae fed on same type of maize cultivar infected with *A. brasilense* and *A. chroococcum* in both years (*P*<0.05; Fig.4). Furthermore, there were significant decreases in RGR, ECD, and ECI, and significant increases in RCR and AD of *M. separata* larvae fed on *Bt* maize in contrast to non-*Bt* maize infected with same type of azoto species within the same CO_2_ level in both years.

**Fig 4.**
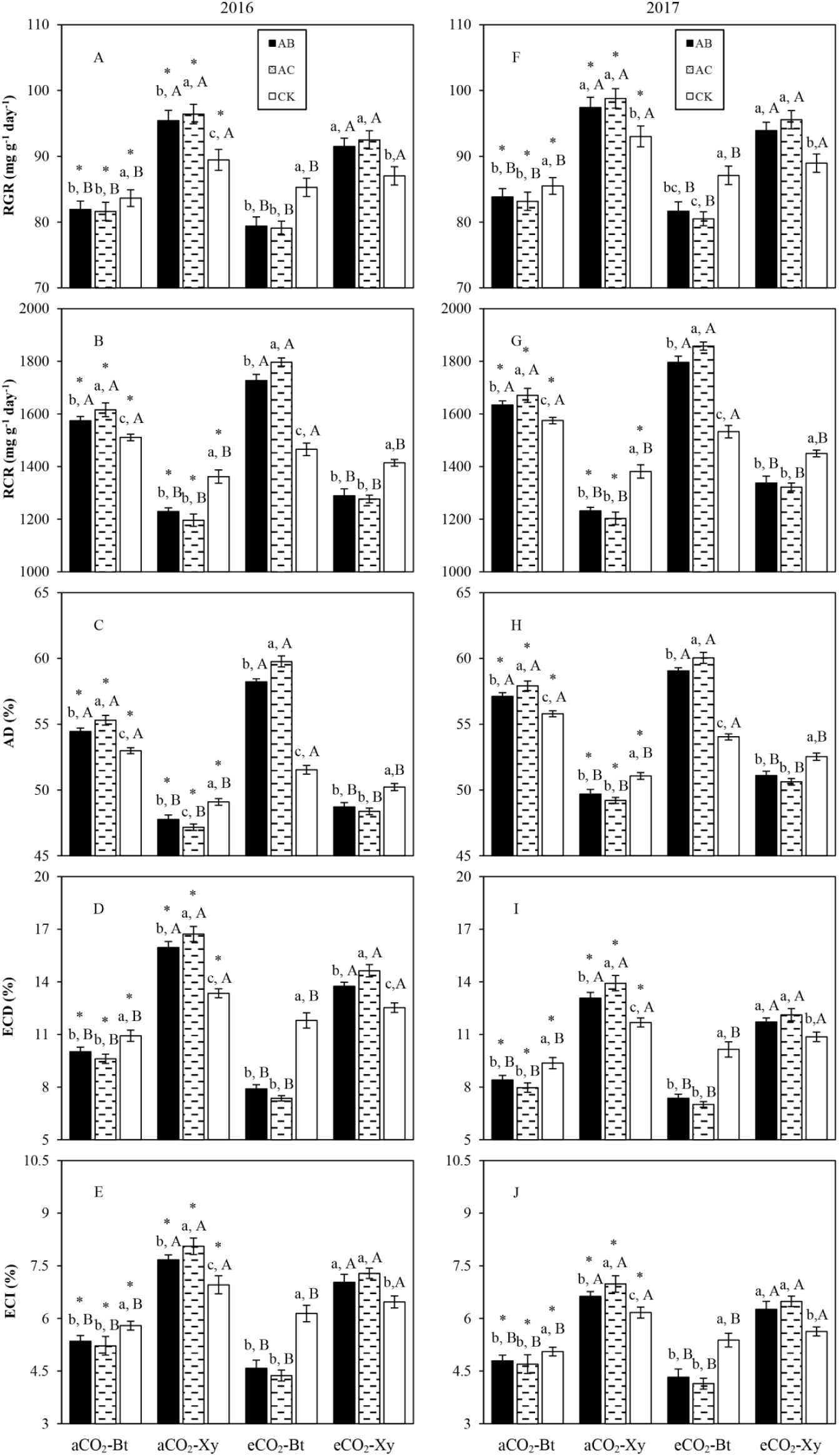
Impacts of the tri-interactions among CO_2_ level, transgenic treatment and azoto infection on the food utilization of *M. separata* from the 3rd to the 6th instar larvae fed on the detached leaves of *Bt* maize (i.e., Bt) and non-*Bt* maize (Xy) during the heading stage, infected with *A. brasilense* (i.e., AB) and *A. chroococcum* (AC) as well buffer solution as the control (CK) under ambient and elevated CO_2_ in 2016 (A–E) and 2017 (F–J). Each value represents the average (+*SE*). Different lowercase and uppercase letters, and * indicated significant difference among three types of azoto infection for same type of maize under same CO_2_ level, between *Bt* maize and non-Bt maize for same type of azoto infection under same CO_2_ level, and between ambient and elevated CO_2_ for same type of maize and azoto infection by the Duncan test at *P*<0.05 respectively.

## Discussion

Insects are sensitive to environmental variations, and environmental stresses can cause changes on their growth, development, fecundity, food utilization and the occurrence and distribution of populations as a result of metabolic rate fluctuation (Bloom et al., 2010). Elevated CO_2_ negatively affected the larval survival, weight, duration, pupation, and adult emergence of cotton bollworm, *Helicoverpa armigera* (Akbar et al., 2016), and reduced the egg laying by Cactus moth *Cactoblastis cactorum* (Stange et al., 1997) and *Achaea Janata* (Rao et al., 2016). In this study, elevated CO_2_ significantly prolonged larval and pupal duration and decreased pupation rate and pupal weight of *M. separata* compared to ambient CO_2_. Relative growth rates (i.e., RGR) of Gypsy moth (*Lymantria dispar*) were reported to be reduced by 30% in larvae fed on *Quercus petraea* exposed to elevated CO_2_ (Hattenschwiler & Schafellner, 2004). Relative consumption rate (i.e., RCR) was significantly higher for *H. armigera* larva fed maize grown at 375 and 750 ppm CO_2_ in contrast to ambient CO_2_ condition, and elevated CO_2_ significantly decreased the efficiency of conversion of ingested food (i.e., ECI), the efficiency of conversion of digested food (i.e., ECD), and the RGR of *H. armigera* larvae compared with ambient CO_2_ (Yin et al., 2010). In this study, elevated CO_2_ significantly increased the RCR (+10.44%) and the approximate digestibility (+5.59%) (i.e., AD), and significantly reduced the RGR (-9.95%), ECD (-16.05%) and ECI (-17.95%) of *M. separata* larvae compared with ambient CO_2_. According to the “Nutrition compensation hypothesis”, elevated CO_2_ can affect the development fitness of herbivores by changing the nutritional components, above and below-ground biomass, and photosynthetic rate of host plants indirectly (Ainsworth & Rogers, 2007; Jackson et al., 2009; Zavala et al., 2013), including increased C/N ratio and decreased nitrogen content etc. Declined growth rate, reproduction, and survival rate were found in the chewing mouthparts insects (e.g. *Helicoverpa armigera*, *Spodoptera exigua*, *Mythimna separata*), and the food consumption of which increased so that they could obtain necessary nutrition to survive (Bottomley et al., 1993; Rogers et al., 2006). Yin *et al.*(2010) reported that elevated CO_2_ increased the food consumption and prolonged the development time of *H. armigera*, which due to the reduced nutritional quality of maize leaves, as a result of reduced nitrogen content and increased C/N ratio. Elevated CO_2_ significantly reduced the food conversion rate and enhanced the food ingestion of *H. armigera*, which attribute to reduced nitrogen content of the cotton, Simian-3 (Chen et al., 2005a; Chen et al., 2005b). Thus, Chen *et al.* (2005a; 2005b) inferred that elevated CO_2_ might be unfavorable to *H. armigera*. Our results in maize system appear to be similar to the study by Chen *et al.* (2005a; 2005b) in a cotton system.

Although the transgenic corn, *Zea mays* L., hybrids expressing the Cry insecticidal protein from *Bacillus thuringiensis* (Bt) were developed to control *Helicoverpa zea, Ostrinia nubilalis, Spodoptera frugiperda* and *Mythimna separata* (Ostlie et al., 1997; Koziel et al., 1993; Armstrong et al., 1995; Jouanin et al., 1998; Lynch et al., 1999), few studies focused on the defense responses of transgenic *cry1Ie* maize to corn armyworm under elevated CO_2_, especially on the growth, development and food utilization of the pest insects. Prutz & Dettner (2005) reported that the transgenic *Bacillus thuringiensis*-maize could result in decreased growth rate and increased mortality, which might attribute to the termination of larval metamorphosis. Most studies showed that adverse effects on life-table parameters of different herbivores were direct by the *Cry* protein (Lawo et al., 2010), which might be due to the interaction of feeding inhibitors and growth inhibitors (e.g. secondary plant substances) (Smith & Fischer, 1983). Effects of elevated CO_2_ on the plant nutrition, metabolism and secondary defense metabolism might adverse for the growth, development and nutrition utilization of herbivores (Akbar et al., 2016). The insects possessed more nutrients to meet their growth needs and prolong the food digestion time in the midgut so that the RCR and AD increased (Reynolds, 1985). In this study, we found that some negative effects of transgenic *cry1Ie* maize (Bt) and Xianyu335 (Xy) grown in elevated CO_2_ on the food utilization indices (including RGR, ECD and ECI) of *M. separata* larvae and some positive effects on the RCR and AD, which indicated that the resistance responses of *Bt* maize might persist under elevated CO_2_, and *M. separata* might ingest more food to get enough nutrition for surviving in limited developmental time under elevated CO_2_. Meanwhile the *Bt* maize and its parental line (Xianyu 335) prolonged their larval life-span and pupal duration, decreased growth rate and increased mortality that might result in lowering of pests’ occurrence. According to the “carbon nutrition balance hypothesis” (Gebauer et al., 1997), elevated CO_2_ would increase the fixed organic matter in plant while increase C-based secondary metabolites and decrease N-based secondary metabolites, thus affecting the insects resistance of plants. Robinson *et al.* (2012) indicated that elevated CO_2_ increased 19% phenols, 22% condensed tannins and 27% flavonoids, while the terpenoids and NBSC decreased by 13% and 16% respectively. Coviella (2002) anticipated that the primary CO_2_ effect on *Bt* toxin production would be due to differences in N concentration within the plant. In a meta-analytical review of 33 studies that simultaneously increased carbon dioxide conditions compared to ambient conditions, Zvereva & Kozlov (2006) showed that nitrogen concentration in plants was reduced under elevated CO_2_, and this decrease was stronger for woody compared to herbaceous plants. If conditions of increased carbon (e.g. elevated CO_2_) allow plants to allocate significantly more resources to condensed tannins and gossypol, then the enzyme composition in the insect herbivore is expected to also change. Similarly, if *Bt* toxin production changes due to elevated CO_2_, then the insect herbivore’s body enzymes should also be changed in this circumstance.

Most of nitrogen, however, is found in the form of nitrogen gas (N_2_) which approximately amounts to 78% in the atmosphere. As plants cannot use this form of nitrogen directly, some microbes can change the nitrogen gas into ammonia. Most free living microbes in soil which can fix nitrogen and whose activities in enhancing the growth of plants are bacteria namely *Azotobacter* sp. and *Azospirillum* sp. These two bacteria are particularly important in maize production system due to their greater nitrogen fixing ability. Azospirillum acquires carbohydrate directly from sieve tube as a resource of carbon which promotes its growth (Olivera et al., 2004). Azospirillum can be used to promote the growth of sprouts under normal and arid conditions (Alejandra et al., 2009). Azospirillum also provides more flexibility to cell wall which enhances the growth (Pereyra et al., 2010) and increases products of wheat in waterless plot of land (Martin et al., 2009). Furthermore, azospirillum had the highest efficiency in nitrogen fixation at the root of sweet corn and it would reach the highest point of nitrogen fixation in the week 4 amounting to 0.20 mgNhr^−1^m^−2^ (Toopakuntho, 2010). Azospirillum can also create auxin, a substance promoting growth of maize, of 53.57 mg/ml (Phookkasem, 2011). Therefore, we used techniques of azotobacter (*A. brasilense* & *A. chroococcum*) inoculation of maize seeds to stimulate plant N uptake to increase in biomass N relative to C under elevated CO_2_, increase *Bt* toxin production for transgenic *cry1Ie* maize and create a substance promoting maize plant growth. In this study, we found that elevated CO_2_ significantly enhanced the rhizosphere soil densities both *A. brasilense* and *A. chroococcum*. We hypothesize that the elevated CO_2_ increased the maize root bifurcation and soil nutrition (e.g. carbohydrates, amino acids and multi-trace elements) for azotobacter to provide the living space and nutrition. Other researchers have also shown positive effects of elevated CO_2_ on the bacterial community in the rhizosphere of maize (Chen, 2012). Moreover, significant advserse effects on the growth, development, reproduction and food utilization of *M. separata* was observed when the host substrate maize was exposed to azotobacter treatments, which might be attributed to azotobacter stimulating plant N uptake to increase *Bt* toxin production for transgenic *cry1Ie* maize and promoting growth of its parental line (Xianyu335).

There was no significant year-to-year variation in our field research data. Therefore, the overall results clearly indicate that increasing CO_2_ had negative effects on *M. separata*. Resistance performance of transgenic *cry1Ie* maize decreased under elevated CO_2_ as shown by decreased RGR, ECD, and ECI. The azotobacter treatments (*A. brasilense* & *A. chroococcum*) had positive effects on improving the effectiveness of *Bt* maize on target Lepidoptera pest management via decreased RGR, ECD, and ECI of *M. separata* that fed on transgenic *cry1Ie* maize and promoting growth of Xianyu 335 via increased RGR, ECD, and ECI of *M. separata*. Under future predicted climate changes (e.g. elevated CO_2_), it is particularly important to understand the field insect resistance traits of resistant crops to target pests. In an environment of accelerated greenhouse effect, *Bt* maize may have decreased resistance performance in the field with inhibiting effect on the development and food utilization of insects. Therefore, we used techniques of azotobacter (*A. brasilense* & *A. chroococcum*) inoculation of maize seeds to stimulate plant N uptake to increase in biomass N relative to C under elevated CO_2_, increase *Bt* toxin production for transgenic *cry1Ie* maize, and create a substance promoting maize growth. This study demonstrates that the use of azotobacter (e.g., *A. brasilense* and *A. chroococcum*) as pest control enhancer especially under elevated CO_2_ is significantly more beneficial in transgenic *Bt* maize system compared to that in non-transgenic system.

## Acknowledgements

Many thanks to Prof. Dr Megha N. Parajulee, Texas A&M University, AgriLife Research and Extension Center, for his help in the design and treatment setup of this experiment, and revising the manuscript.

## Data availability

All data are included within the manuscript.

